# Topological Pharmacokinetics: Reading the Shape of Drug Disposition from Data

**DOI:** 10.64898/2026.05.13.724751

**Authors:** Hong-Can Ren, Ya-Xin Gu

**Affiliations:** Department of DMPK, Genescience Pharmaceuticals Co., Ltd. Research and Development, Shanghai, China

## Abstract

Pharmacokinetic analysis has spent half a century compressing drug concentration–time curves into scalar summaries—AUC, C_max_, clearance—discarding the shape information that encodes mechanistic fingerprints of the underlying physiology. We introduce Topological Pharmacokinetics (TPK), a framework that reads the shape of pharmacokinetic trajectories directly from data without prior commitment to a compartmental model. TPK uses delay embedding to reconstruct the pharmacokinetic attractor from the concentration–time curve, and persistent homology to extract its topological invariants—connected components and loops—as a Pharmacokinetic Topological Invariant (PTI) vector. We validate TPK across three levels: linear systems (negative control), nonlinear saturable elimination (detection of the N_PTP +1 rule and a nonlinear diagnostic triad), and endogenous circadian rhythms (contrastive detection of rhythmic interference via Dev specificity and Decouple Collapse). The PTI vector provides a model-agnostic shape fingerprint that, in simulation, demonstrates the diagnostic potential of shape-based analysis; validation on experimental data is required to assess whether this potential generalizes to real pharmacokinetic data. All findings are demonstrated as proof of concept on simulated data; validation on experimentally measured concentration–time curves is the essential next step.

## INTRODUCTION

Pharmacokinetics characterizes what the body does to a drug through two established approaches.[1,2] Non-compartmental analysis reduces the concentration–time curve to scalar summaries—AUC, Cmax, half-life—which are robust and regulatory-accepted, but discard shape by construction. Compartmental modeling restores fidelity by fitting sums of exponentials, but only captures shapes belonging to the pre-selected function family. When data deviate—a secondary peak from enterohepatic recirculation,[3,4] a bowed elimination phase from saturable clearance,[5,6] a periodic baseline from a circadian rhythm[7,8]—these deviations are relegated to the residual error term, modeled as independent Gaussian noise. This constitutes a systematic shape blind spot: information that reveals mechanism is treated as nuisance variance.

The blind spot is consequential. Enterohepatic recirculation produces a secondary peak that, absent a pre-specified biliary compartment, is absorbed entirely into the residuals; in the language of TPK, this is a persistent H1 generator—a loop in the reconstructed phase space.

Saturable elimination—whether due to capacity-limited metabolism, saturable protein binding, or nonlinear transport—systematically bends the elimination phase away from log-linearity, producing a structured arc in the residuals; TPK reads this arc as a curvature transition point (N_PTP), detectable without knowing Km. Endogenous circadian or ultradian rhythms superimpose periodic oscillations onto drug concentration profiles; the standard practice of subtracting a constant baseline flattens these oscillations, manufacturing spurious secondary features or masking genuine pharmacodynamic feedback.[8,9] TPK detects such oscillations as a specific, invariant shape distortion (Dev) that survives superposition with the exogenous drug, and further reveals the rhythm’s loss of dynamical autonomy through a Decouple Collapse.

Topological Data Analysis offers a fundamentally different approach. Persistent homology—the core engine of TDA—tracks topological features (connected components, loops) as they appear, persist, and disappear across a multiscale filtration of data, extracting robust shape descriptors without parametric assumptions.[10,11] Long-lived features correspond to genuine geometric structures; short-lived ones are attributed to noise—a distinction guaranteed by the stability theorem of persistent homology.[12,13] In the life sciences, TDA has been applied to characterize neuronal branching,[14] gene regulatory network dynamics, single-cell transcriptomic architecture,[15] and protein conformational landscapes.[16,17] Pharmacokinetics is not merely another application domain; it is a domain in which the geometry of the data is the direct imprint of the physiology that generated it, and in which the ability to read shape without a model addresses a blind spot that has persisted for half a century.

This paper introduces Topological Pharmacokinetics (TPK). A drug’s concentration–time trajectory is the observable trace of a dynamical system—absorption, distribution, metabolism, and excretion—evolving on a low-dimensional manifold M. The state vector x(t) includes not only plasma concentration but also tissue drug levels, enzyme occupancy, transporter activities, and physiological rhythms. We observe only a one-dimensional projection C(t) = h(x(t)). Delay embedding, justified by Takens’ theorem,[18] reconstructs from C(t) alone a point cloud diffeomorphic to M.[19] Persistent homology then extracts topological invariants—connected components, loops, and their persistence across scales—as a Pharmacokinetic Topological Invariant (PTI) vector: a fixed-length, model-agnostic shape fingerprint. Because the PTI vector is computed via a fixed pipeline with no model selection step, it provides a standardized, machine-readable feature set—an AI-ready representation suitable for large-scale pharmacokinetic data integration.

We validate TPK across three levels of escalating complexity. First, benchmark validation on linear pharmacokinetic systems establishes the negative control: TPK correctly reports the absence of cyclic structure where none exists. Second, detection of nonlinear saturable elimination reveals an empirically invariant topological signature of nonlinear elimination under the conditions tested—the N_PTP +1 rule—alongside a nonlinear diagnostic triad of independent shape-based criteria. Third, a contrastive strategy for detecting endogenous circadian rhythms without a clean baseline identifies Dev as a rhythm-specific sentinel and reveals a topological coupling (Decouple Collapse) that is detectable from shape data alone, though its mechanistic interpretation requires further experimental validation. Collectively, these investigations establish TPK as a principled, model-free complement to existing methodology—one whose diagnostic potential, demonstrated here in simulation, warrants systematic evaluation on clinical data.[20,21]

## METHODS

### Theoretical Framework

#### Pharmacokinetic systems as dynamical flows

The ADME system involves a multitude of interacting entities—drug molecules in plasma, tissue-bound drug, enzyme occupancy states, transporter activities, metabolite concentrations, and physiological rhythms—defining a high-dimensional state vector **x**(*t*)evolving on a smooth, low-dimensional manifold ℳ, constrained by mass balance and physiological regulation: 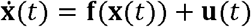, where **f** encodes the intrinsic pharmacokinetic dynamics and **u**(*t*) represents the externally imposed dosing regimen.[1] The manifold ℳ is the set of all states the system actually visits; its dimension reflects the true degrees of freedom of the system. We do not observe **x**(*t*) directly. The plasma concentration is obtained via the observation function *h* that projects the manifold onto a one-dimensional time series:

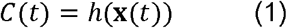

where *C*(*t*) is the measured plasma drug concentration at time *t*, **x**(*t*) ∈ ℳ is the full pharmacokinetic state vector whose coordinates include tissue drug levels, enzyme occupancy, transporter states, and physiological rhythms, and *h*: ℳ → ℝ is the observation function that projects the high-dimensional trajectory onto a single dimension. Information is lost in this projection: features clearly separated on ℳ may overlap in the scalar record, and geometric structures such as loops may be flattened beyond recognition. The central hypothesis is that despite this loss, the concentration–time series retains sufficient structure to recover the essential topological features of ℳ—guaranteed under generic conditions by Takens’ embedding theorem.[18,19]

We acknowledge that clinical PK data deviate from the idealized conditions of Takens’ theorem in two respects. First, the theorem assumes a continuous, noise-free trajectory—a mathematical ideal unavailable in any experimental setting. Pharmacokinetic time series are discretely sampled and contaminated by measurement error. The stability theorem of persistent homology provides partial protection, ensuring that small perturbations produce proportionally small changes in the persistence diagram.[12] Second, the embedding dimension must exceed twice the intrinsic manifold dimension, a quantity that remains unknown for any real pharmacokinetic system. Under these conditions, the reconstructed point cloud is a practical approximation whose quality depends jointly on sampling density, signal-to-noise ratio, and the chosen embedding parameters.

#### Normalization and delay embedding

Prior to the construction of delay vectors, the concentration time series is normalized by its maximum observed value to eliminate the confounding effects of dose magnitude and ensure the scale invariance of topological features:

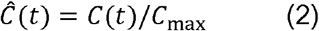

where *C*_max_ = max_t_ *C*(*t*). Delay vectors are then constructed from the normalized time series as:

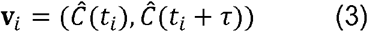

where *t*_*i*_ is the *i*-th sampling time and *τ* is the time delay. For the uniformly sampled pharmacokinetic profiles in this study, *τ* is set to 5 sampling intervals, corresponding to 1.0 h when the time step is 0.2 h, or approximately 0.4 h when the time step is 0.08 h. A two-dimensional embedding (*m* = 2) is employed because a single delay coordinate is sufficient to unfold the pharmacokinetic attractor and to capture prominent cyclic or curved structures, while facilitating visualization and computational efficiency. The set of all such delay vectors 𝒫 = {**v**_*i*_} constitutes the reconstructed point cloud. Takens’ theorem guarantees that this embedding preserves the topological features of the original manifold under generic conditions.

For clinically sparse data (typically 5–15 points per subject),[6,7] naive delay embedding into ℝ^*m*^ with *m* ≥ 3 produces too few points to resolve topology reliably. We propose model-assisted probabilistic embedding: a population PK model is fit to the full dataset to obtain population parameter estimates and inter-individual variability.[20,22] For a subject with sparse observations, Bayesian posterior estimation yields a posterior predictive distribution over plausible dense trajectories.[23] An ensemble of *K* dense trajectories is simulated from this posterior, and topological stability is tested via pairwise bottleneck distances of their persistence diagrams.[24] If the maximum bottleneck distance falls below a stability threshold (set as the 95th percentile of null distances from permuted data), the topology is deemed robust to posterior uncertainty, and the median trajectory’s embedding is selected. If unstable, the method transparently reports that the available data are insufficient to resolve the topology.

A terminological clarification is warranted. The population PK model is used exclusively as a densification engine—it provides a principled means of interpolating between sparse observations without itself being the object of topological interrogation. The PTI vector is computed from the reconstructed point cloud, not from the estimated parameters of the interpolating model. In the dense-sampling limit, model-assisted embedding is bypassed entirely and the pipeline operates in a fully nonparametric mode. In the sparse-sampling regime, the method depends on a population PK model for densification but does not use that model to interpret the resulting shape features. Moreover, the topological stability test provides a safeguard: when the posterior ensemble fails to converge on a consistent topology, the method transparently reports uncertainty. TPK is best characterized as “model-assisted but not model-dependent.” All experiments reported in this paper used dense sampling (300 points), bypassing model-assisted embedding entirely; the model-assisted sparse-sampling extension described above has not been empirically evaluated in the present work.

#### Persistent homology

Persistent homology tracks topological features across a multiscale Vietoris–Rips filtration of the embedded point cloud 𝒫. For the one-dimensional character of the concentration–time curve, zero-dimensional persistent homology (*H*_0_) is equivalent to a prominence analysis of the time series: the prominence of a peak, defined as the vertical distance from the peak to the highest contour line that separates it from a higher peak, equals exactly the persistence of the corresponding connected component when the filtration parameter is interpreted as the concentration level. We therefore compute *H*_0_ features efficiently via peak detection using scipy.signal.find_peaks with a prominence threshold of 0.05 × *C*_max_ (applied prior to normalization, on the raw concentration scale). The number of detected peaks, *N*_sig_ (*H*_0_), and their prominence values, normalized by *C*_max_ to yield the persistence *P* (*H*_0_), serve as the *H*_0_-derived components of the PTI vector.

One-dimensional persistent homology (*H*_1_) is computed using Ripser[25] on the normalized two-dimensional delay vectors. A loop is considered significant if its persistence exceeds a fixed threshold *ε*_noise_ = 0.015 in the normalized space. This empirical cutoff was selected by applying the TPK pipeline to all six linear pharmacokinetic benchmark systems (IV and Oral, one- and two-compartment) and verifying that no H1 feature exceeded this value; the highest observed H1 persistence in any linear system was 0.008, providing a margin of approximately 1.9× below the threshold. This calibration procedure is circular in the sense that the threshold was derived from the same benchmark systems used to validate the negative control result; the permutation-based procedure described in Supplementary S2 provides an independent, non-circular validation but was not applied to the scenarios reported in the main text. Under the dense-sampling, low-noise conditions used throughout this study, this cutoff reliably separates genuine cyclic structures from noise; its generalization to different signal-to-noise ratios or sampling densities requires the permutation-based procedure described in Supplementary S2.

In pharmacokinetic terms, *H*_0_ generators correspond to temporally separated phases of the concentration–time trajectory—for example, the absorption phase forming one cluster and the elimination phase forming another. *H*_1_ generators are the topological signature of cyclic or strongly curved processes: enterohepatic recirculation, where a secondary concentration peak creates a closed orbit in the delay-embedded phase space;[3,4] saturable elimination, where the transition from zero-order to first-order kinetics may produce a bowed trajectory;[5,6] and endogenous circadian rhythms, where a periodic physiological driver imprints a stable cycle on the data.[7,8]

#### The PTI vector

From the normalized point cloud 𝒫 and its persistence diagram PD_*k*_(𝒫), we extract the Pharmacokinetic Topological Invariant (PTI) vector. Its components are organized into two classes: Class A (strict topological invariants) and Class B (geometry-infused descriptors computed after topological structure is confirmed).

The number of significant *H*_1_ generators counts the *H*_1_ features whose persistence exceeds the noise threshold:

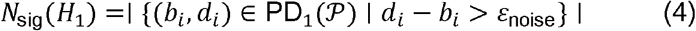

where ε_noise_ = 0.015 for the normalized phase space used throughout this work. In a purely monotonic, linear elimination system, *N*_sig_ (*H*_1_) = 0. A value of 1 indicates a single dominant cyclic process; values ≥ 2 suggest multiple nested or sequential feedback mechanisms.

The maximum *H*_1_ persistence is *P*_max_(*H*_1)_ = max {*d*_*i*_ − *b*_*i*_ | (*b*_*i*_, *d*_*i*_) ∈ PD_1_(𝒫)}, the persistence of the most robust cycle in the system. If PD_1_(𝒫) is empty, *P*_max_ (*H*_1_) = 0. The maximum H_0_ persistence is *P*_max_ (*H*_0_) = max_*i*_ {prom_*i*_}/ *C*_max_, reflecting the maximal separation between distinct phases of the pharmacokinetic trajectory.

The topological entropy quantifies the structural complexity of the profile using the Shannon entropy of the normalized prominence distribution of all detected peaks:

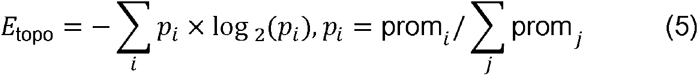

A system dominated by a single peak has *E*_topo_ ≈ 0; a system with multiple peaks of comparable prominence has higher *E*_topo_.

The number of phase transition points, *N*_PTP_, is obtained by locating curvature peaks along the embedded trajectory. The local curvature of the two-dimensional trajectory *γ* (*t*) = (*Ĉ* (*t*), *Ĉ* (*t* + *τ*)) is computed as:

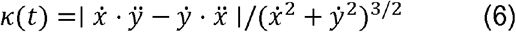

where *x* (*t*) = *Ĉ* (*t*), *y* (*t*) = *Ĉ* (*t* + *τ*), and derivatives are approximated by central finite differences. Peaks in *κ* (*t*) exceeding a prominence threshold of max (10^−5^,0.05 × max_*t*_ *κ* (*t*)) are identified as phase transition points; their count defines *N*_PTP_. In the current toolkit version, only *N*_PTP_ is reported; the maximum curvature value κ_max_ itself is reserved for future extensions.

The shape distortion index quantifies the departure of the elimination phase from ideal first-order behavior as the maximum perpendicular deviation from the secant line connecting the endpoints of the elimination segment in the delay-embedded space:

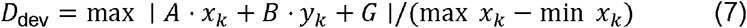

where (*x*_*k*_,*y*_*k*_) = (*Ĉ* (*t*_*k*_), *Ĉ* (*t*_*k*_ + *τ*)) are the delay-embedded points of the elimination phase (starting at the last significant peak), and *A, B, G* are the coefficients of the straight line connecting the first and last points of that segment. The denominator normalizes by the *x*-axis span, rendering *D*_dev_ scale-invariant. A value near zero confirms linear elimination; a large value flags systematic deviation from exponential decay.

The saddle decoupling index quantifies the degree of separation between two topologically distinct kinetic events:

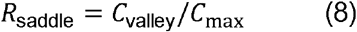

where *C*_valley_ is the minimum concentration between the two most prominent peaks on the raw (un-normalized) concentration–time curve. A value approaching 0 indicates cleanly decoupled processes (e.g., an immediate-release pulse followed by a well-separated delayed-release pulse); a value approaching 1 indicates merging of phases.

The inter-peak interval Δ*t* is defined as the median time difference between all consecutively detected peaks on the concentration–time curve. When a significant *H*_1_ loop is present, this interval generally reflects the cycle period.

The topological polarity index integrates shape significance with exposure intensity:

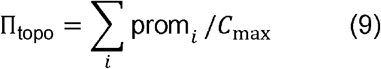

where the sum runs over the prominences of all detected peaks. A profile with high exposure but a simple shape yields a small Π_topo_; a profile with complex shape embedded in a moderate concentration range yields a larger Π_topo_.

The topological nonlinearity index is a unitless global measure:

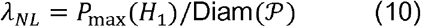

where Diam (𝒫) = max_*i,j*_ ∥ **v**_*i*_ - **v**_*j* ∥_ is the diameter of the embedded point cloud—the maximum pairwise Euclidean distance between any two delay vectors. Normalizing by the overall scale renders *λ*_*NL*_ comparable across dose levels and individuals. A system with *λ*_*NL*_ → 0 is topologically linear; a system with *λ*_*NL* ≫_ 0 is topologically nonlinear. In the present work, *λ*_*NL*_ is reported for completeness, but its detailed empirical evaluation is deferred to future studies.

### Benchmark Validation: Linear Pharmacokinetics

We applied TPK to six simulated systems: IV 1-compartment, IV 2-compartment, Oral 1-compartment, Oral 2-compartment, Oral dual-site absorption (DSA), and Oral enterohepatic recirculation (EHC). All models received a dose of 1000 mg. The IV 1-compartment model used V = 10 L and k = 0.2 h^− 1^. The Oral 1-compartment model used ka = 1.2 h^− 1^ and k = 0.2 h^− 1^. DSA was constructed by superimposing a delayed absorption component (lag time 4 h, ka = 1.5 h^− 1^, relative fraction 0.5) onto the Oral 1-compartment base. EHC was constructed by adding a gallbladder-emptying pulse at 12 h (ka = 0.8 h^− 1^, relative fraction 0.4) onto the same base. All trajectories were densely sampled (300 points, 0–24 h). Delay embedding used m = 2 and τ = 5 sampling steps (approximately 0.4 h). Vietoris–Rips persistent homology was applied to the normalized point clouds.

### Shape Discovery: Nonlinear Pharmacokinetics

Twenty-four scenarios fully crossed route (IV, Oral), compartments (1, 2), elimination (linear, Michaelis-Menten with Vmax = 10 mg/h and Km = 5 mg/L), and dose (200, 1000, 4000 mg).[5,6] Under this parameterization, the Low dose already approaches Km; Mid and High extend deeply into the saturated regime. All simulations used dense sampling with additive Gaussian noise (σ = 0.5).

### Penetrating the Baseline: Endogenous Rhythms

Three concentration–time trajectories spanning 120 h (0.1-h resolution) were generated: a 24-hour circadian oscillation given by C(t) = 10 + 4·cos(2πt/24) (amplitude 4% of the exogenous peak concentration), a one-compartment IV bolus (Cmax = 100, tdose = 12 h, k = 0.2 h^− 1^), and their linear superposition. All signals were analyzed without added measurement noise to isolate the intrinsic topological signatures of the underlying dynamics. We employed a contrastive strategy: comparing the PTI vectors of the pure exogenous drug (Exo) and the mixed signal (Mix) rather than attempting physical separation. The difference ΔPTI constitutes the topological footprint of the endogenous rhythm.

## RESULTS

### Benchmark Validation: Topological Consistency with Linear Pharmacokinetics

Figure 1 presents the full TPK pipeline output for all six benchmark systems——concentration–time curves, persistence diagrams, and delay-embedded phase spaces. Table 1 reports the corresponding PTI parameters. Across all four linear systems, N_sig(H1) = 0—no spurious cyclic signals (Table 1). IV 1-compartment exhibited a perfectly contractible trajectory (D_dev = 0, N_PTP = 0). IV 2-compartment showed measurable curvature (D_dev = 0.060, N_PTP = 1) without generating a closed loop—TPK correctly distinguishes multi-compartment curvature from cyclic dynamics. Oral 1-compartment produced the characteristic teardrop-shaped trajectory but no persistent H1 cycle: oral absorption, when linear, does not produce false-positive cyclic signals. Oral 2-compartment generated two curvature transition points without H1 generators.

**Table 1.** PTI parameters for benchmark validation (six systems).

| Model | N_H0 | N_sig(H1) | Dev | N_PTP | $\Delta t$ (h) | R_saddle | E_topo | $\Pi_{\text{topo}}$ |
| --- | --- | --- | --- | --- | --- | --- | --- | --- |
| IV 1-Comp | 1 | 0 | 0 | 0 | 0 | — | 0 | 1.00 |
| IV 2-Comp | 1 | 0 | 0.060 | 1 | 0 | — | 0 | 1.00 |
| Oral 1-Comp | 1 | 0 | 0.032 | 1 | 0 | — | 0 | 0.99 |
| Oral 2-Comp | 1 | 0 | 0.060 | 2 | 0 | — | 0 | 0.90 |
| Oral DSA | 2 | 2 | 0.031 | 4 | 3.29 | 0.685 | 0.672 | 1.18 |
| Oral EHC | 2 | 1 | 0.035 | 4 | 12.20 | 0.157 | 0.811 | 1.22 |

**Figure 1.**
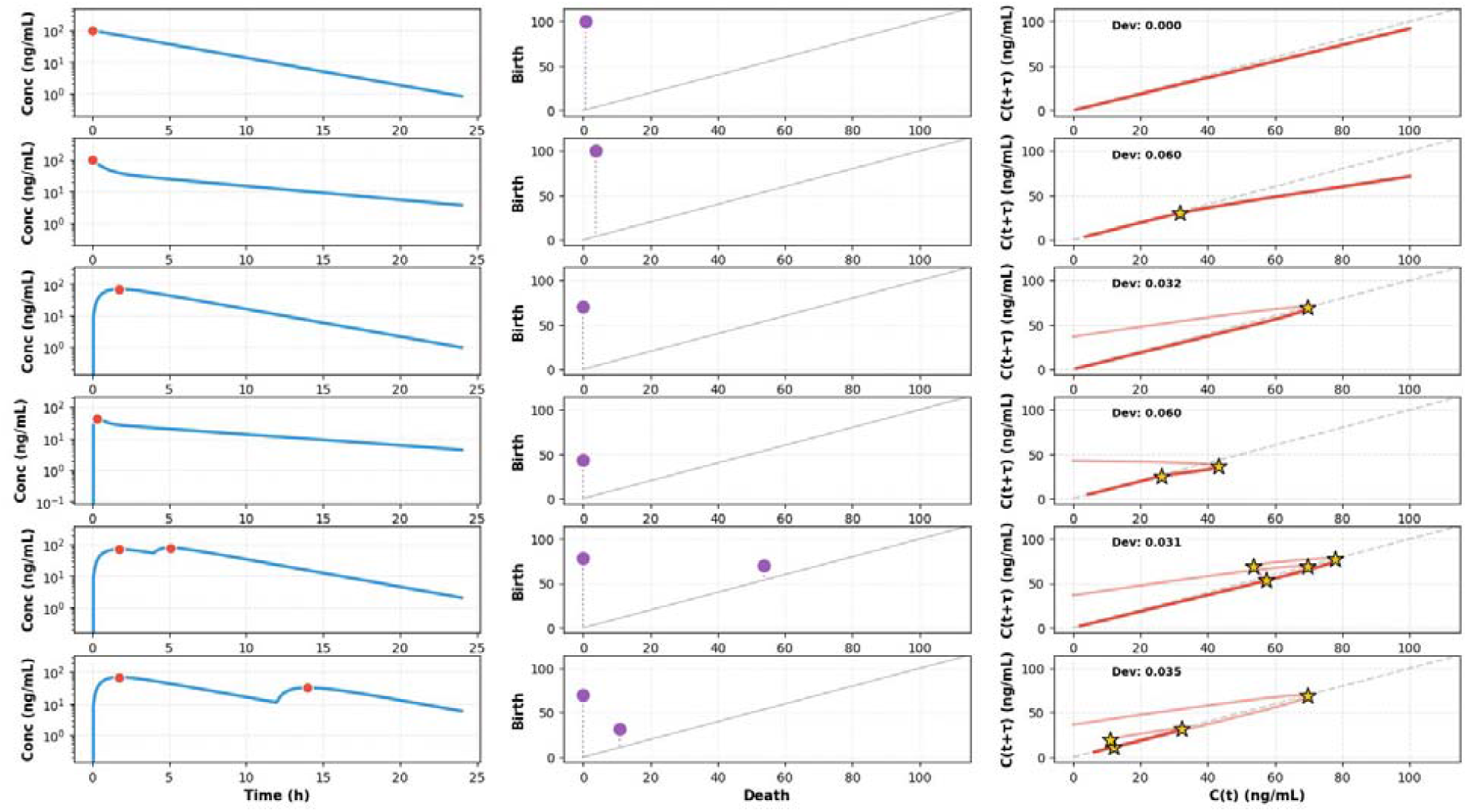
Topological Pharmacokinetics applied to six benchmark systems. Each row shows one pharmacokinetic model (top to bottom: IV 1-Comp, IV 2-Comp, Oral 1-Comp, Oral 2-Comp, Oral DSA, Oral EHC). Left column: concentration–time curves (semi-log scale) with phase transition points (red dots). Middle column: H0 persistence diagrams (Birth vs. Death). Right column: delay-embedded phase space with Dev values. Only Oral DSA (row 5) and Oral EHC (row 6) generate H1 generators (N_sig(H1) ≥ 1), confirming the negative control: linear PK and oral absorption do not produce false cyclic signals.

Oral DSA detected two significant H1 generators (N_sig(H1) = 2, E_topo = 0.672, Δt = 3.29 h, R_saddle = 0.685), the topological signature of concurrent, partially coupled absorption events—the first peak has not fully resolved before the second begins.[4] Oral EHC detected one H1 generator with a substantially longer inter-peak interval (Δt = 12.20 h) and near-complete decoupling (R_saddle = 0.157)—almost four times the DSA interval, consistent with the physiological time scale of biliary excretion and gallbladder emptying.[3,4] The contrast between DSA and EHC—both producing two peaks in the concentration–time profile—establishes TPK’s diagnostic hierarchy. At Level 1, D_dev and N_PTP identify multi-compartment curvature present in IV 2-Comp and Oral 2-Comp. At Level 2, N_sig(H1) ≥ 1 confirms genuine cyclic structure, present only in DSA and EHC. At Level 3, R_saddle distinguishes fully decoupled physiological cycles (EHC, 0.16) from coupled parallel processes (DSA, 0.68). At Level 4, Δt provides mechanistic constraints—short Δt suggests parallel absorption, long Δt suggests sequential physiological cycling. No single conventional pharmacokinetic parameter provides comparable stratified diagnosis.[1,2]

### Shape Discovery: Topological Detection of Nonlinear Pharmacokinetics

Figure 2 displays the concentration–time profiles for all 24 scenarios. Table 2 reports the PTI parameters at the Mid dose level for representative configurations. Across all 24 scenarios, N_sig(H1) = 0. This is not a detection failure but a precise topological distinction that completes the diagnostic hierarchy: nonlinear elimination produces curvature, not cycles—in contrast to the genuine H1 generators from EHC and DSA observed above.[5,6] Linear systems exhibited shape rigidity: D_dev invariant across dose levels (Table 2). Figure 3 confirms this visually: in the linear panels, dose-normalized trajectories from different dose levels collapse onto a single curve in the phase space. In contrast, every nonlinear system exhibited manifold distortion—dose-dependent shape variation in at least one PTI parameter. The most robust indicator was N_PTP: every nonlinear system exhibited exactly one more curvature transition point than its linear counterpart (IV 1-Comp: 0→1; IV 2-Comp: 1→2; Oral 1-Comp: 1→2; Oral 2-Comp: 2→3). The N_PTP +1 rule, as observed across the 24 scenarios in our factorial design, constitutes an empirically invariant topological signature of nonlinear elimination under the conditions tested. To our knowledge, this is the first model-free geometric invariant of pharmacokinetic nonlinearity reported in the literature. The additional curvature transition point corresponds to the transition from the saturated regime to the first-order regime as concentration declines.

**Table 2.** PTI parameters for representative nonlinear scenarios (Mid dose, 1000 mg).

| Route | Comp | Elim | Dev | N_PTP | E_topo | $\Pi_{\text{topo}}$ |
| --- | --- | --- | --- | --- | --- | --- |
| IV | 1 | Linear | 0 | 0 | 0 | 1.00 |
| IV | 1 | Nonlinear | 0.003 | 1 | 0 | 1.00 |
| IV | 2 | Linear | 0.064 | 1 | 0 | 1.00 |
| IV | 2 | Nonlinear | 0.081 | 2 | 0 | 1.00 |
| Oral | 1 | Linear | 0.025 | 1 | 0 | 1.00 |
| Oral | 1 | Nonlinear | 0.005 | 2 | 0 | 1.00 |
| Oral | 2 | Linear | 0.050 | 2 | 0 | 1.00 |
| Oral | 2 | Nonlinear | 0.022 | 3 | 0 | 1.00 |

**Figure 2.**
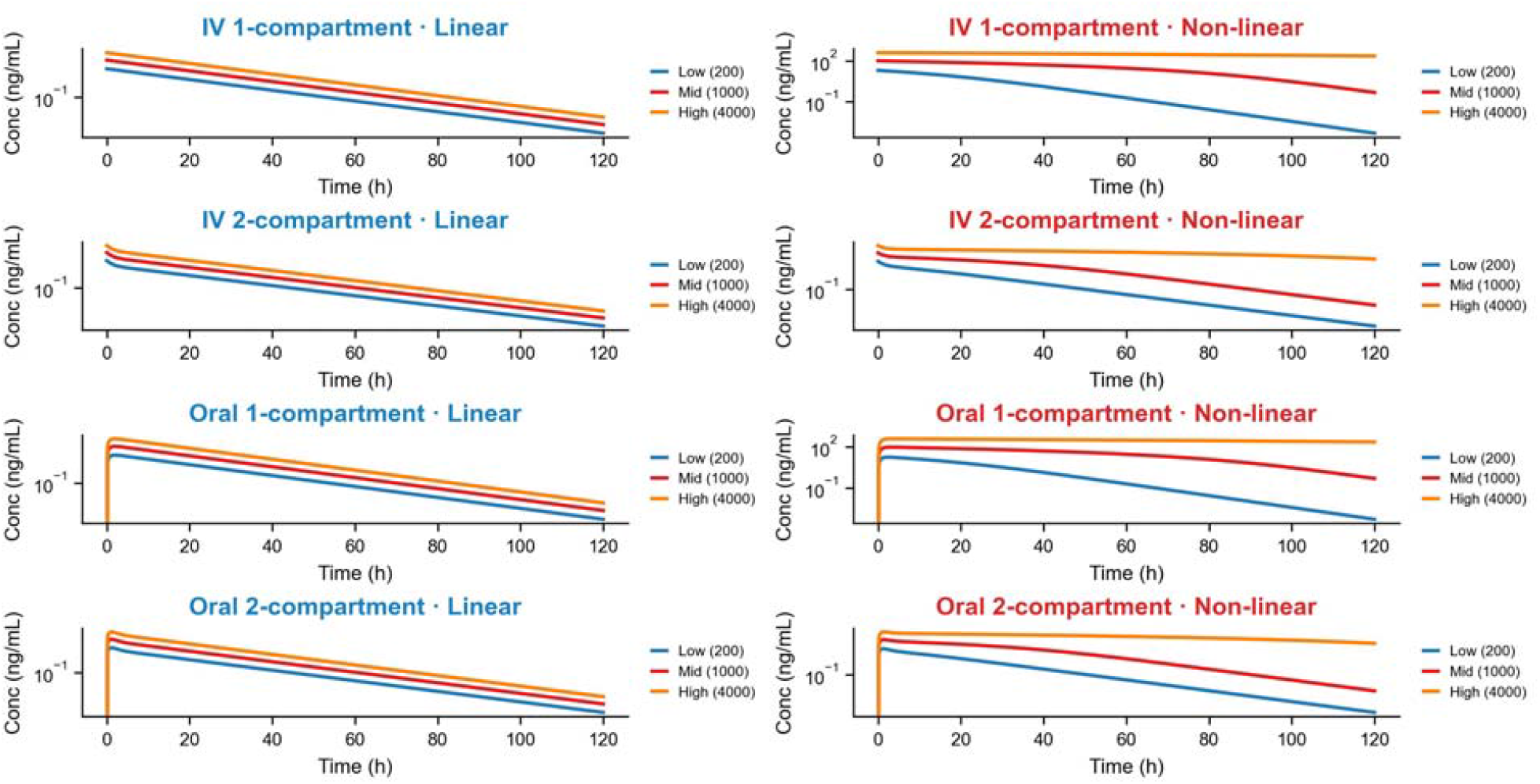
Simulated concentration–time profiles for linear (left, blue titles) and nonlinear Michaelis–Menten elimination (right, red titles) across four route×compartment combinations at three dose levels: Low (200 mg, blue), Mid (1000 mg, red), High (4000 mg, orange). Linear systems exhibit dose-proportional parallel curves (shape rigidity); nonlinear systems exhibit dose-disproportionate profiles with bowed elimination phases. See Methods for simulation parameters.

**Figure 3.**
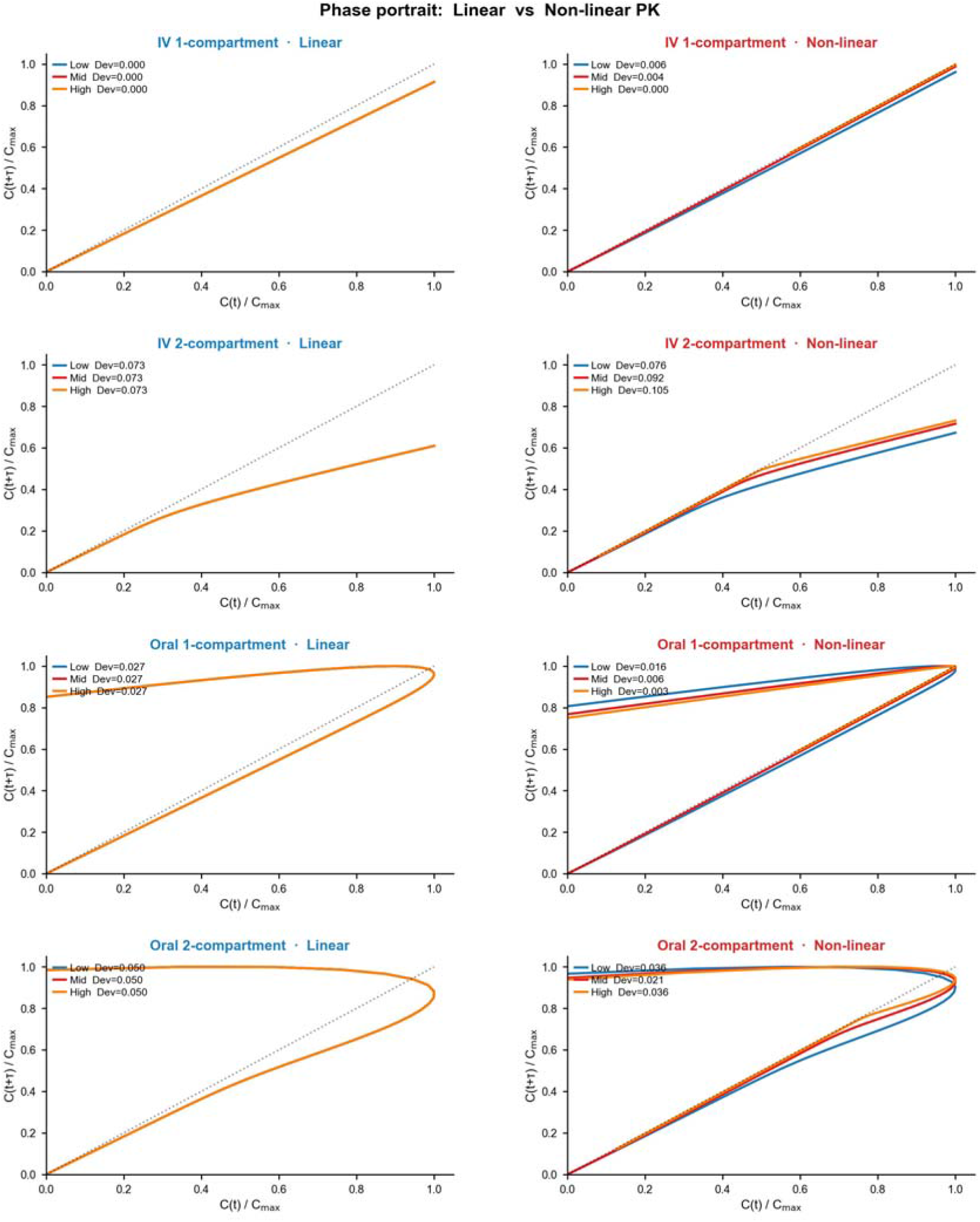
Phase portraits (C(t)/C_max vs. C(t+τ)/C_max) comparing linear (left, blue) and nonlinear (right, red) PK across four route×compartment combinations at three dose levels. In linear systems, Dev is dose-invariant (overlapping curves); in nonlinear systems, Dev varies systematically with dose. The N_PTP +1 rule manifests as an additional curvature inflection in each nonlinear panel. Note: the dose-dependence of Dev in the IV 2-Comp nonlinear panel (range 0.076–0.105) is subtle visually; precise values are provided in Table 2.

In contrast, the nonlinear panels of Figure 3 show dose-dependent spread of the phase-space trajectories, reflecting the manifold distortion induced by saturable elimination. D_dev in IV 2-Comp nonlinear increased monotonically with dose (0.066→0.080→0.091). The biphasic distribution compounded with saturable elimination produces progressive deviation from the linear reference. In contrast, IV 1-Comp nonlinear D_dev decreased with dose (0.0047→0.0031→1.6×10^−5^) — saturation straightens the trajectory: near-zero-order elimination produces a trajectory closer to a straight line than the curved first-order trajectory. In oral nonlinear systems, Π_topo dropped at the highest dose (0.42 and 0.88), reflecting topological flattening of the attractor as saturable elimination prolongs exposure. Together, N_PTP elevation, Dev dose-dependence, and Π_topo decline constitute a nonlinear diagnostic triad. A system satisfying any two of these three criteria can be identified as topologically nonlinear without specifying a parametric model.

### Penetrating the Baseline: Topological Detection of Endogenous Rhythms

Figure 4 shows the concentration–time curves for the three signals. Figure 5 presents the complete TPK analysis——PK curves, persistence diagrams, phase spaces, and curvature κ(t)——for the Endo, Exo, and Mix signals in separate rows. Table 3 reports the PTI comparison. The mixed signal inherited the rhythm’s H1 generator (N_sig(H1) = 1 vs. Exo = 0). Dev was the most striking finding: Endo = 1.410, Exo = 0, Mix = 1.410—numerically identical to the endogenous signal (Table 3). This equality is mathematically expected: Dev measures the departure of the elimination phase from log-linearity in the delay-embedded space, and when a linear exogenous drug is superimposed on an endogenous oscillation, the elimination arc of the composite trajectory inherits the shape distortion of the oscillatory component unchanged. The result is therefore a structural property of the linear superposition, not an independently calibrated empirical finding. Its practical significance is nonetheless real: a non-zero Dev in a system where linear elimination is expected constitutes a diagnostic flag for an unmodeled periodic baseline, detectable without baseline measurement, pre-dose sampling, or a parametric model of the rhythm.

**Table 3.** PTI comparison for endogenous rhythm, exogenous drug, and mixed signal. □ Shift = Mix − Exo. For N_PTP, the additive expectation (Endo + Exo = 17 + 0 = 17) is exceeded by 10 emergent curvature transition points; the Shift value of +27 reflects the absolute change from Exo alone, not the excess over the linear sum.

| Parameter | Endo | Exo | Mix | Shift(Mix-Exo) |
| --- | --- | --- | --- | --- |
| N_sig(H1) | 1 | 0 | 1 | +1 |
| Dev | 1.410 | 0 | 1.410 | +1.410 |
| N_PTP | 17 | 0 | 27 | +27□ |
| R_saddle | 0.385 | — | 0.052 | — |
| E_topo | 2.415 | 0 | 1.066 | +1.066 |

**Figure 4.**
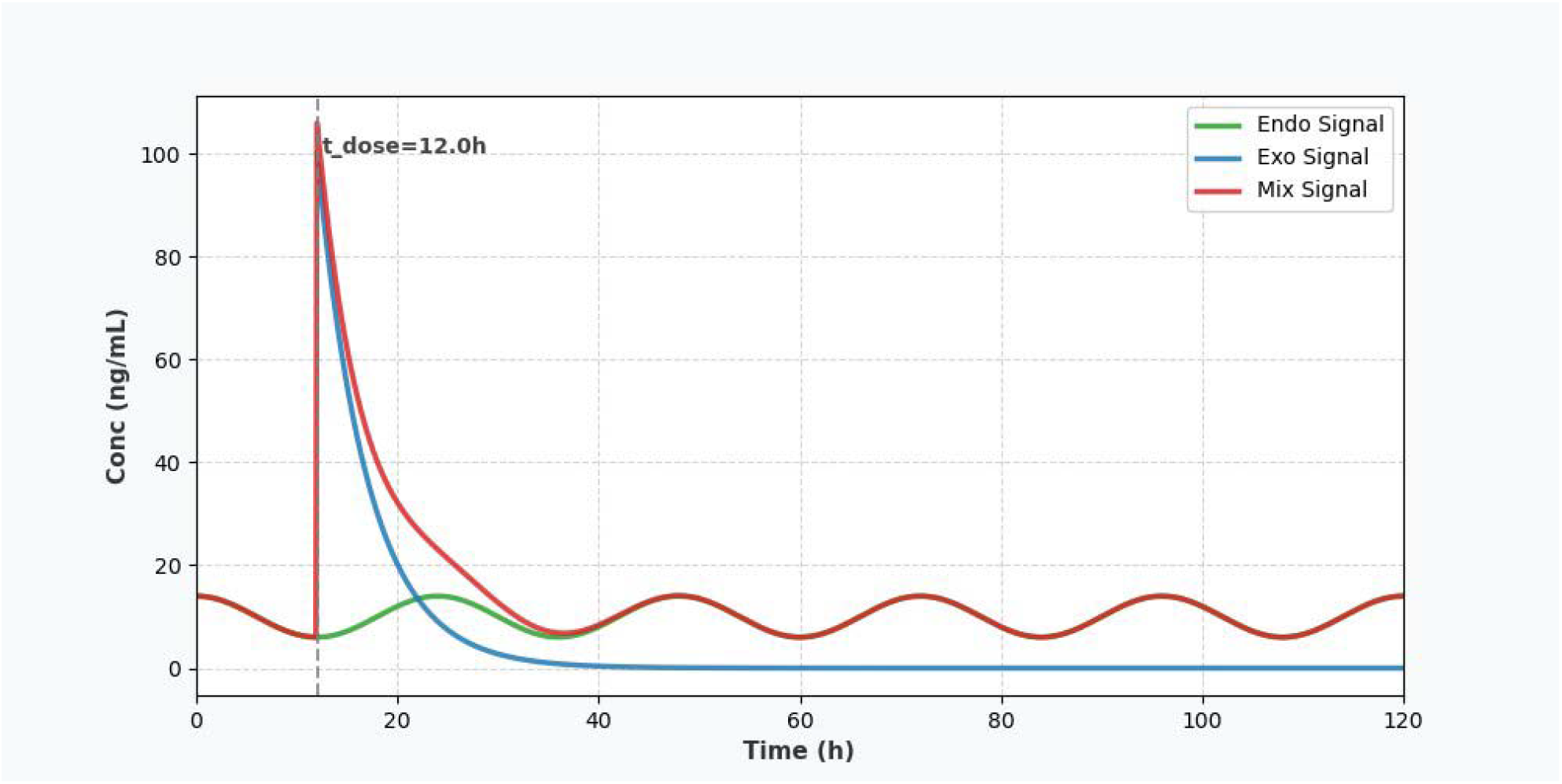
Simulated concentration–time profiles for the three signals in the endogenous rhythm experiment (120 h; t_dose = 12 h). Green: Endogenous circadian oscillation (Endo; C(t) = 10 + 4·cos(2πt/24)). Blue: Exogenous IV bolus (Exo; one-compartment, k = 0.2 h^− 1^, C_max = 100 ng/mL at t = 12 h). Red: Linear superposition of both signals (Mix). The Mix signal inherits the oscillatory baseline superimposed on exponential decay.

**Figure 5.**
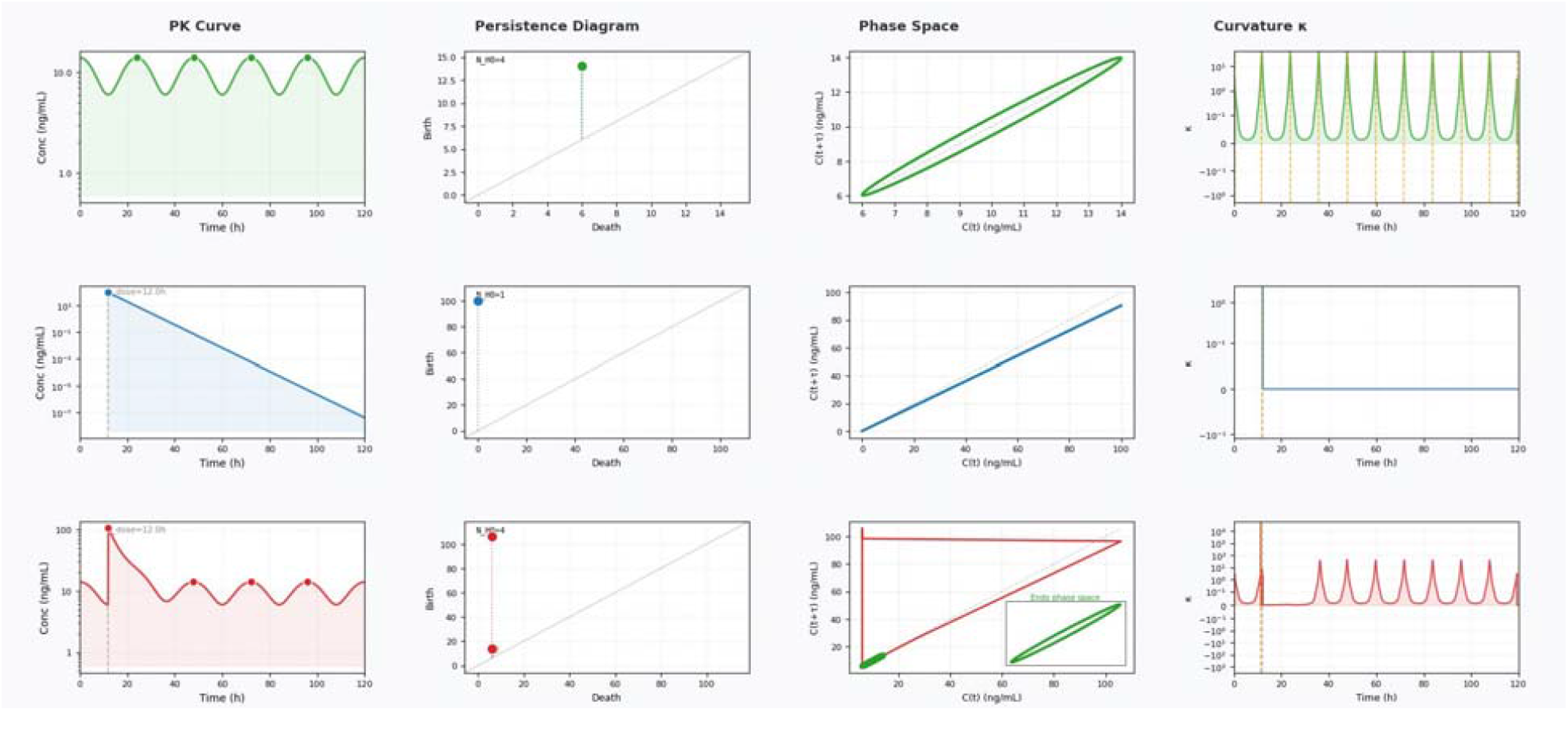
Full TPK analysis of the three endogenous rhythm signals. Rows (top to bottom): Endo (green), Exo (blue), Mix (red). Columns (left to right): PK curve with dose time marked, H0 persistence diagram, delay-embedded phase space, curvature κ vs. time. Endo shows an elliptical H1 loop and periodic κ spikes. Exo shows a monotone diagonal trajectory (no loop) and near-zero post-dose curvature. Mix inherits the H1 loop (N_sig(H1) = 1), N_PTP = 27 (10 emergent curvature transitions above the additive expectation of Endo + Exo = 17), and R_saddle compression from 0.385 (Endo) to 0.052 (Mix). The inset in the Mix phase space panel shows the Endo attractor overlaid.

N_PTP(Mix) = 27 exceeded the additive expectation of 17 by 10 curvature transition points. These extra curvature transitions are not contributed by either signal alone; they are emergent features of rhythm–drug interaction. When the weak circadian oscillation modulates the steep exponential decay, the composite trajectory acquires new inflection points. The saddle decoupling index dropped from 0.385 (Endo) to 0.052 (Mix). The substantial decrease in R_saddle indicates that the dominant topological features of the mixed trajectory are more tightly coupled than those of the endogenous rhythm alone. Pharmacokinetically, the rhythm’s peaks and troughs—which were distinct and temporally separated in the endogenous state—become bound to the drug’s elimination arc in the mixed signal, losing their character as features of an independent oscillation. This topological coupling is a geometric fact of the superposition, detected without ambiguity. Whether this coupling reflects a genuine pharmacodynamic interaction—in which the drug alters the rhythm’s underlying regulatory machinery—or is simply the arithmetic consequence of a linear sum of two independent time series, cannot be determined from shape data alone. TPK’s contribution is the detection and quantification of the coupling; mechanistic interpretation requires experimental designs that go beyond the scope of the present work.

The contrastive strategy—comparing PTI vectors rather than attempting physical separation—reveals four independent signatures: N_sig(H1) inheritance, Dev specificity, N_PTP excess, and Decouple Collapse. Critically, the presence or absence of an H1 generator cleanly separates nonlinear elimination (Beta_1 = 0 in all 24 scenarios) from endogenous rhythms (Beta_1 = 1, inherited by the mixed signal). Nonlinear elimination bends the trajectory (curvature without cycles); endogenous rhythms loop it (curvature with cycles). This completes TPK’s diagnostic hierarchy (Table 4).

**Table 4.**
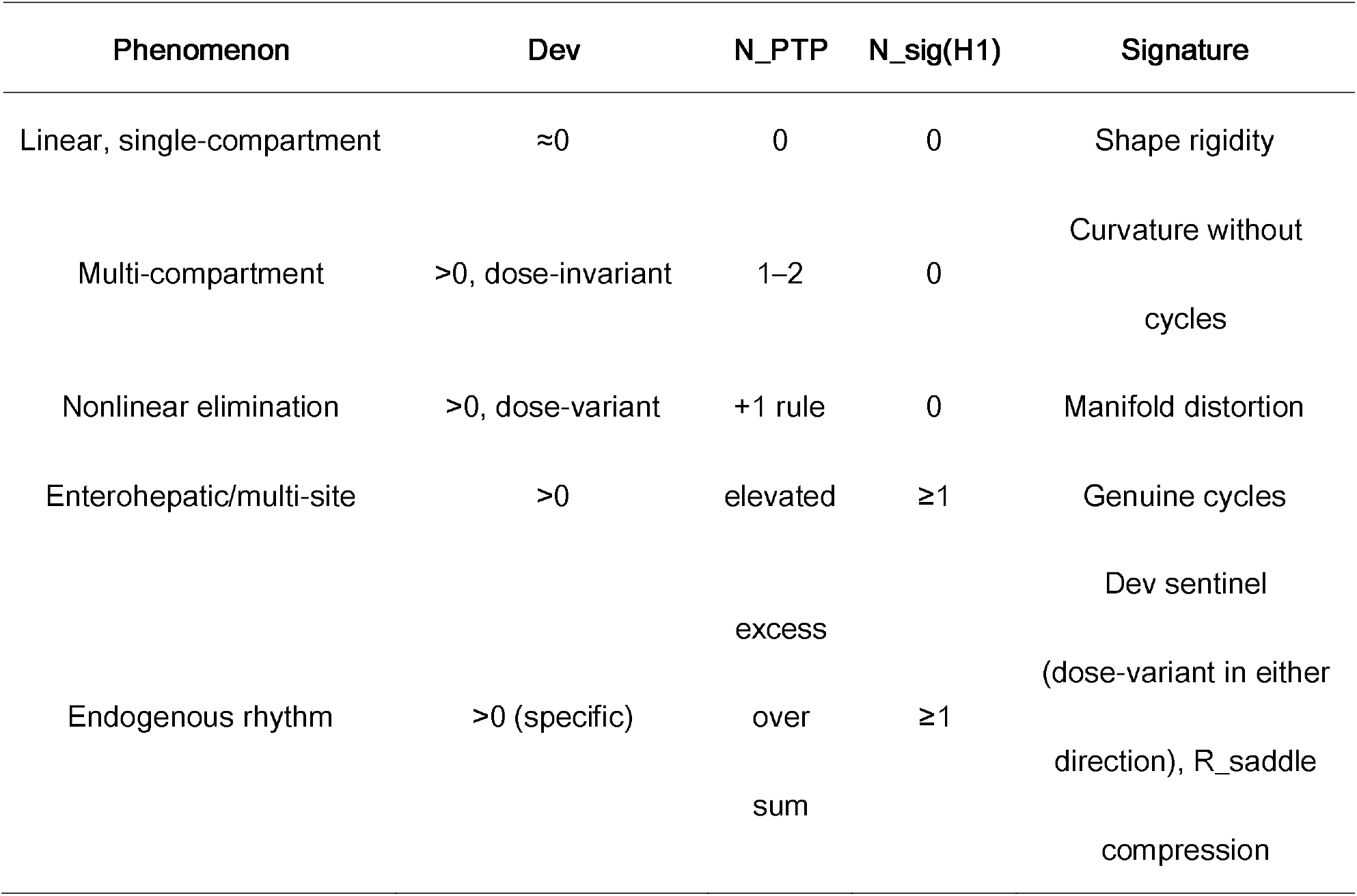
TPK diagnostic hierarchy.

## DISCUSSION

The investigations reported here constitute a progressive validation of a single central claim: that the shape of a pharmacokinetic trajectory, read through persistent homology, carries diagnostic information that conventional scalar analysis cannot access. The evidence stands on three independent pillars: negative control in linear systems; detection of nonlinear elimination through the N_PTP +1 rule and nonlinear diagnostic triad; and contrastive detection of endogenous rhythms through Dev specificity and Decouple Collapse.

### Theoretical significance

TPK’s defining capability is recovering high-dimensional dynamical information from a low-dimensional observable. The concentration–time curve is a one-dimensional projection of the full ADME manifold. Delay embedding reconstructs a point cloud diffeomorphic to that manifold; persistent homology reads its topological invariants. What is recovered is not the full geometry—that would require observing all state variables—but its topological essence: the loops, connected components, and persistent structures that survive projection. Persistent homology distills the invariant topological core of the trajectory; geometric information above the topological level is not retained, but the topological invariants themselves are stable and reproducible. This is a mathematically principled complement to conventional statistical compression, operating on shape rather than magnitude.[18,19,26]

### AI-ready pharmacokinetics

The PTI vector transforms the concentration–time curve from a model-dependent, irregularly sampled time series into a fixed-length vector in a geometrically meaningful shape space. In this space, similar pharmacokinetic dynamics cluster together regardless of dose amplitude or route of administration. This is the prerequisite for large-scale machine learning on pharmacokinetic data. Just as convolutional neural networks rely on translation-invariant feature maps in image space, future AI models for pharmacokinetics can rely on model-agnostic, dose-normalized shape features in PTI space. TPK provides the eye; PTI space provides the visual cortex.

### A common language for shape

TPK establishes a common vocabulary for pharmacokinetic shape. Terms like “H1 generator,” “persistence,” “shape rigidity,” “manifold distortion,” and “Decouple Collapse” are not metaphors. They are mathematically defined quantities with invariant geometric meaning, computable identically on any concentration–time curve. This common language makes possible comparisons that were previously impossible: between drugs, between studies, between healthy and diseased states—providing the foundation for a cumulative science of pharmacokinetic shape.

Limitations. Current limitations include the uncharacterized boundaries of sparse sampling,[27] the undeveloped statistical inference infrastructure for persistence diagrams,[10,11] and the challenge of separating dosing-driven periodicity from endogenous cycles in multi-dose regimens. A significant barrier to regulatory or clinical adoption is the absence of hypothesis tests, confidence intervals, and sample size calculations for PTI parameters—the mathematical statistics of persistent homology are still maturing, and the permutation-based noise threshold, while practical, has not been validated against known-distribution scenarios. The Vietoris-Rips filtration scales quadratically in the number of data points, constraining application to very long time series without subsampling strategies. The generality of the N_PTP +1 rule across a wider range of Km values, sampling schedules, and noise levels remains to be systematically characterized. The choice of the embedding delay parameter τ (set to 5 sampling intervals throughout) was not subjected to sensitivity analysis; changes in τ can alter N_PTP counts, Dev values, and H1 persistence, and the sensitivity of PTI parameters to this free parameter requires systematic characterization before the method can be applied in new settings. All findings reported here are based on simulated data, which provides ground truth essential for theoretical validation; the next critical step is validation on experimentally measured concentration–time curves. The diagnostic hierarchy was demonstrated on clearly separated, idealized simulations; its performance on ambiguous or overlapping cases remains to be evaluated.

### Extension toward Topological Pharmacology

The logic of TPK extends with full symmetry to pharmacodynamic data. A pharmacodynamic time series—blood pressure, T-cell activation, tumor volume—can be analyzed by the same delay-embedding and persistent-homology pipeline without assuming a specific PK/PD model.[28,29] The same analytical pipeline can be applied to effect–time curves, capturing tolerance loops,[30,31] rebound dynamics,[32] and chronopharmacodynamic modulation.[33] The development of Topological Pharmacodynamics (TPD) is the immediate next step in our research program. When PK and PD PTI vectors are placed in a common shape feature space, the result is a unified Topological Pharmacology (TP): a single geometric language describing a drug’s fate and effect.[34,35] The ultimate horizon extends to quantitative systems pharmacology,[36,37,38] where persistent homology may extract the functional skeleton of complex biological systems—feedback loops, modular separations, persistent cycles—that remains invariant under parameter uncertainty.[39,40] The eye that TPK gives to the computer begins with a single curve; its destination is the geometry of life.

Pharmacokinetics has spent half a century asking *how much*—how much drug is in the blood, how large is the exposure, how fast is it cleared. Topological Pharmacokinetics asks *what shape*. In the curve of drug concentration over time, there is geometry: loops where physiology cycles, bends where kinetics saturate, structures where endogenous rhythms persist. This geometry has always been there, encoded in the data, but invisible to tools designed to compress curves into numbers. We have shown, in simulation, that this geometry can be read. The blind spot, for now, has been mapped; it has not yet been fully illuminated.

## STUDY HIGHLIGHTS

### What is the current knowledge on the topic?

Pharmacokinetic analysis reduces concentration–time curves to scalar summaries (AUC, Cmax) or fits pre-specified compartmental models, systematically discarding shape information that encodes mechanistic fingerprints of processes such as enterohepatic recirculation, nonlinear elimination, and endogenous rhythms.

### What question did this study address?

Can the shape of a pharmacokinetic trajectory be read directly from data—without pre-committing to a compartmental model—to detect nonlinearity, cyclic processes, and endogenous baseline interference?

### What does this study add to our knowledge?

We introduce Topological Pharmacokinetics (TPK), combining delay embedding and persistent homology to extract a Pharmacokinetic Topological Invariant (PTI) vector. In simulation, TPK reveals an empirically invariant N_PTP +1 rule for nonlinear elimination (empirically invariant across the 24 scenarios tested), a nonlinear diagnostic triad, and a contrastive strategy that detects endogenous rhythms through Dev specificity and topological coupling. Validation on experimentally measured concentration–time curves is required before these findings can be generalized.

### How might this change drug discovery, development, and/or therapeutics?

TPK provides a model-agnostic, AI-ready shape fingerprint for pharmacokinetic data. It can serve as an automated sentinel for nonlinearity and rhythmic interference in early drug development, and its extension to pharmacodynamics (TPD) opens the path toward a unified topological pharmacology.

## AUTHOR CONTRIBUTIONS

Hon-Can Ren conceived and directed the research, developed the theoretical framework of Topological Pharmacokinetics, and led manuscript writing and revision. Ya-Xin Gu implemented the Python PTI extraction pipeline, generated all simulation data, and contributed to the design of key topological parameters and visualization outputs. All authors reviewed and approved the final manuscript.

## AI ASSISTANCE STATEMENT

**DeepSeek (深度求索, DeepSeek-R1)** assisted with theoretical development, manuscript drafting and revision, equation formulation, and parameter system design. **Anthropic Claude** (**version 3.5 Sonnet**) provided independent peer review prior to submission. All core ideas, scientific interpretations, and conclusions are the intellectual contribution of the human authors, who critically reviewed and verified all AI-generated content and take full responsibility for the accuracy and integrity of the published work.

## DATA ACCESSIBILITY STATEMENT

All simulation code used to generate the results in this paper, including the PTI vector extraction pipeline implemented in Python using Ripser[25] and GUDHI libraries, is available at [The code is available at https://github.com/yxgu2353/TopoPK]. The complete simulated datasets and PTI output tables are included in the Supplementary Information.

## SUPPLEMENTARY INFORMATION

Additional supporting information, including the complete 24-scenario PTI output table (Table S2), the L3 parameter directory (Table S1), detailed algorithmic workflows for model-assisted probabilistic embedding, and supplementary methods for permutation-based noise threshold determination, accompanies this article online.

