## Supplemental Tables and Methods for "Topological Pharmacokinetics: Reading the Shape of Drug Disposition from Data"

**Table S1. Pharmacokinetic Topological Invariant (PTI) Parameter Tiered Classification System**

| **Tier** | **Symbol** | **Parameter Name** | **Definition** | **Pharmacokinetic Interpretation** |
| --- | --- | --- | --- | --- |
| L1 | $N_{\text{sig}}(H_{1})$ | Number of Significant H1 Generators | $\mid\{(b_{i},d_{i})\in\text{PD}_{1}\mathcal{(P)\mid}\mathcal{l}_{i}>\epsilon_{\text{noise}}\}\mid$ | Count of robust cyclic processes (enterohepatic recirculation, dual-site absorption, endogenous rhythms) |
| L1 | $P_{\text{max}}(H_{1})$ | Maximum H1 Persistence | $\max\{\mathcal{l}_{i}\mid(b_{i},d_{i})\in\text{PD}_{1}\mathcal{(P)\}}$ | Scale and robustness of the dominant cycle |
| L1 | $P_{\text{max}}(H_{0})$ | Maximum H0 Persistence | $\max\{\mathcal{l}_{i}\mid(b_{i},d_{i})\in\text{PD}_{0}\mathcal{(P)\}}$ | Maximal separation between trajectory phases (e.g., absorption vs. elimination) |
| L1 | $E_{\text{topo}}$ | Topological Entropy | $-\sum_{i} p_{i}\log_{2}(p_{i})$, $p_{i}=\mathcal{l}_{i}/\sum_{j} \mathcal{l}_{j}$ | Structural complexity of the attractor; distinguishes simple from multi-mechanism profiles |
| L2 | $D_{\text{dev}}$ | Shape Distortion Index | $d_{F}(\gamma_{\text{elim}},L)$, Fréchet distance from linear reference | Quantifies departure from log-linear elimination; zero = linear, positive = curvature |
| L2 | $\kappa_{\text{max}}$ | Maximum Phase Space Curvature | $\max_{t}\parallel\dot{\gamma}(t)\times\ddot{\gamma}(t)\parallel/\parallel\dot{\gamma}(t)\parallel^{3}$ | Sharpest kinetic transition (e.g., absorption-to-elimination switch) |
| L2 | $N_{\text{PTP}}$ | Number of Phase Transition Points | Number of curvature peaks along embedded trajectory | Number of distinct kinetic regime switches; +1 rule for nonlinearity |
| L2 | $A_{\text{loop}}$ | H1 Generator Loop Area | Shoelace area of PCA-projected cycle points | Volume of drug participating in recirculation; separates large from small cycles |
| L2 | $\Pi_{\text{topo}}$ | Topological Polarity Index | $\sum_{i} \mathcal{l}_{i}/C_{\text{max}}$ | Shape-aware exposure metric: significance vs. intensity |
| L3 | $\Delta t(H_{1})$ | Inter-Peak Interval | $\mid t_{\text{peak},2}-t_{\text{peak},1}\mid$ for significant H1 | Physiological transit time estimate (gallbladder emptying, intestinal reabsorption) |
| L3 | $R_{\text{saddle}}$ | Saddle Decoupling Index | $C_{\text{valley}}/C_{\text{peak, low}}$ | Degree of process separation; 0 = fully decoupled, 1 = merged |
| L3 | $T_{\text{birth, main}}$ | Main Peak Birth Time | Time at maximum topological prominence | Primary pharmacokinetic event onset |
| L3 | $C_{\text{birth, main}}$ | Main Peak Birth Level | Concentration at maximum topological prominence | Primary event concentration |
| L3 | $T_{\text{birth, sub}}$ | Secondary Peak Birth Time | Time at second-highest topological prominence | Secondary event onset (if present) |
| L3 | $C_{\text{birth, sub}}$ | Secondary Peak Birth Level | Concentration at second-highest topological prominence | Secondary event concentration (if present) |

**Table S2. Complete PTI Output for 24 Pharmacokinetic Scenarios (Chapter 4)**

| **Route** | **Comp** | **Elim** | **Dose** | $\boldsymbol{D}_{\text{dev}}$ | $\boldsymbol{N}_{\text{PTP}}$ | $\boldsymbol{N}_{\text{sig}}\boldsymbol{(}\boldsymbol{H}_{\boldsymbol{1}}\boldsymbol{)}$ | $\boldsymbol{E}_{\text{topo}}$ | $\boldsymbol{\Pi}_{\text{topo}}$ | $\boldsymbol{\lambda}_{\boldsymbol{NL}}$ |
| --- | --- | --- | --- | --- | --- | --- | --- | --- | --- |
| IV | 1 | Linear | 200 | $9.1\times{10}^{-11}$ | 0 | 0 | 0 | 1.00 | 0 |
| IV | 1 | Linear | 1000 | $9.1\times{10}^{-11}$ | 0 | 0 | 0 | 1.00 | 0 |
| IV | 1 | Linear | 4000 | $9.1\times{10}^{-11}$ | 0 | 0 | 0 | 1.00 | 0 |
| IV | 1 | Nonlinear | 200 | 0.0047 | 1 | 0 | 0 | 1.00 | 0 |
| IV | 1 | Nonlinear | 1000 | 0.0031 | 1 | 0 | 0 | 1.00 | 0 |
| IV | 1 | Nonlinear | 4000 | $1.6\times{10}^{-5}$ | 1 | 0 | 0 | 1.00 | 0 |
| IV | 2 | Linear | 200 | 0.0638 | 1 | 0 | 0 | 1.00 | 0 |
| IV | 2 | Linear | 1000 | 0.0638 | 1 | 0 | 0 | 1.00 | 0 |
| IV | 2 | Linear | 4000 | 0.0638 | 1 | 0 | 0 | 1.00 | 0 |
| IV | 2 | Nonlinear | 200 | 0.0662 | 2 | 0 | 0 | 1.00 | 0 |
| IV | 2 | Nonlinear | 1000 | 0.0805 | 2 | 0 | 0 | 1.00 | 0 |
| IV | 2 | Nonlinear | 4000 | 0.0910 | 2 | 0 | 0 | 1.00 | 0 |
| Oral | 1 | Linear | 200 | 0.0248 | 1 | 0 | 0 | 1.00 | 0 |
| Oral | 1 | Linear | 1000 | 0.0248 | 1 | 0 | 0 | 1.00 | 0 |
| Oral | 1 | Linear | 4000 | 0.0248 | 1 | 0 | 0 | 1.00 | 0 |
| Oral | 1 | Nonlinear | 200 | 0.0142 | 2 | 0 | 0 | 1.00 | 0 |
| Oral | 1 | Nonlinear | 1000 | 0.0049 | 2 | 0 | 0 | 1.00 | 0 |
| Oral | 1 | Nonlinear | 4000 | 0.0026 | 2 | 0 | 0 | 0.42 | 0 |
| Oral | 2 | Linear | 200 | 0.0495 | 2 | 0 | 0 | 1.00 | 0 |
| Oral | 2 | Linear | 1000 | 0.0495 | 2 | 0 | 0 | 1.00 | 0 |
| Oral | 2 | Linear | 4000 | 0.0495 | 2 | 0 | 0 | 1.00 | 0 |
| Oral | 2 | Nonlinear | 200 | 0.0356 | 3 | 0 | 0 | 1.00 | 0 |
| Oral | 2 | Nonlinear | 1000 | 0.0218 | 3 | 0 | 0 | 1.00 | 0 |
| Oral | 2 | Nonlinear | 4000 | 0.0277 | 3 | 0 | 0 | 0.88 | 0 |

*All nonlinear scenarios used Michaelis-Menten elimination with* $V_{\max}=10$ *mg/h and* $K_{m}=5$ *mg/L.* $\lambda_{NL}$ *is reported for completeness; its detailed empirical evaluation is deferred to future studies.*

**Table S3. L3 Interpretation Parameters for Benchmark Validation (Chapter 3)**

| **Model** | $\boldsymbol{T}_{\text{birth, main}}$ **(h)** | $\boldsymbol{C}_{\text{birth, main}}$ | $\boldsymbol{P}_{\text{main}}$ | $\boldsymbol{T}_{\text{birth, sub}}$ **(h)** | $\boldsymbol{C}_{\text{birth, sub}}$ | $\boldsymbol{R}_{\text{pers}}$ | $\boldsymbol{\Delta}\boldsymbol{t}$ **(h)** |
| --- | --- | --- | --- | --- | --- | --- | --- |
| IV 1-Comp | 0 | 100.00 | 1.000 | — | — | — | — |
| IV 2-Comp | 0 | 100.00 | 1.000 | — | — | — | — |
| Oral 1-Comp | 1.766 | 69.877 | 0.986 | — | — | — | — |
| Oral 2-Comp | 0.321 | 43.193 | 0.898 | — | — | — | — |
| Oral DSA | 5.057 | 78.248 | 0.974 | 1.766 | 69.877 | 0.214 | 3.291 |
| Oral EHC | 1.766 | 69.877 | 0.917 | 13.967 | 32.277 | 0.333 | 12.201 |

$P_{\text{main}}$*: persistence of the main peak normalized to* $C_{\max}$*.* $R_{\text{pers}}$*: ratio of secondary peak persistence to main peak persistence.* $\Delta t$*: inter-peak interval.*

**Supplementary Methods S1. Model-Assisted Probabilistic Embedding — Detailed Algorithmic Workflow**

The model-assisted probabilistic embedding procedure operates as follows:

**Step 1.** Fit a population PK model (e.g., a standard one- or two-compartment model with first-order elimination) to the full dataset using NONMEM, Monolix, or Stan, obtaining population parameter estimates (fixed effects) and inter-individual variability (IIV) parameters.

**Step 2.** For a subject with sparse observations $\mathbf{C}_{\text{obs}}=(C(t_{1}),\ldots,C(t_{n}))$ where $n$ is typically 5–15, compute the Bayesian posterior distribution of the individual parameters $\boldsymbol{\theta}_{i}$ given $\mathbf{C}_{\text{obs}}$ using Markov chain Monte Carlo (MCMC) or importance sampling.

**Step 3.** Draw $K$ samples from this posterior: $\boldsymbol{\theta}_{i}^{\left( 1 \right)},\ldots,\boldsymbol{\theta}_{i}^{\left( K \right)}$. Throughout this work, $K=100$ was used.

**Step 4.** For each sample $\boldsymbol{\theta}_{i}^{\left( k \right)}$, simulate a dense concentration–time trajectory at a fine temporal grid (200–500 points spanning the dosing interval) using the same structural model.

**Step 5.** Apply delay embedding ($m=2$, $\tau$ as specified in the main text) to each of the $K$ dense trajectories, producing an ensemble of point clouds $\mathcal{P}^{\left( 1 \right)},\ldots,\mathcal{P}^{\left( K \right)}$.

**Step 6.** Compute persistence diagrams $\text{PD}(\mathcal{P}^{\left( k \right)})$ for each point cloud via Ripser, and measure pairwise bottleneck distances $d_{B}(\text{PD}(\mathcal{P}^{\left( k \right)}),\text{PD}(\mathcal{P}^{\left( k^{'} \right)}))$ for all $k\neq k^{'}$.

**Step 7.** If $\max_{k,k^{'}}d_{B}<\varepsilon_{\text{stability}}$, where $\varepsilon_{\text{stability}}$ is set adaptively as the 95th percentile of bottleneck distances from 1,000 random permutations of the sparse time series, the topological structure is deemed robust to posterior uncertainty. The median trajectory's embedding (selected by minimizing the sum of squared bottleneck distances to all other embeddings) is used for downstream PTI extraction. If $\max_{k,k^{'}}d_{B}\geq\varepsilon_{\text{stability}}$, the method reports "topologically unstable—insufficient sampling to resolve the shape of this pharmacokinetic trajectory," providing direct feedback to experimental design.

**Supplementary Methods S2. Noise Threshold Determination via Permutation Testing**

The noise threshold $\epsilon_{\text{noise}}$ separates genuine topological features from those attributable to sampling noise. It is determined as follows:

**Step 1.** Given the original concentration–time curve $C(t_{1}),\ldots,C(t_{N})$, generate $M=1,000$ permuted time series by randomly shuffling the temporal order of the concentration values. Each permutation destroys the temporal structure (and hence any genuine topological features) while preserving the marginal distribution of concentrations.

**Step 2.** For each permuted time series, apply the full TPK pipeline (delay embedding, persistent homology) to obtain a null persistence diagram $\text{PD}^{\left( \text{null} , m \right)}$.

**Step 3.** Pool all persistence values $\left\{ \mathcal{l}_{i}^{\left( m \right)} \right\}$ from all null diagrams across all permutations.

**Step 4.** Set $\epsilon_{\text{noise}}$ as the 95th quantile of this pooled null distribution. Features in the original persistence diagram with $\mathcal{l}_{i}>\epsilon_{\text{noise}}$ are deemed statistically significant at $\alpha=0.05$.

**Supplementary Methods S3. Fréchet Distance Computation for** $\boldsymbol{D}_{\text{dev}}$

The Fréchet distance between two parametrized curves $\gamma_{1}:[0,1]\to\mathbb{R}^{m}$ and $\gamma_{2}:[0,1]\to\mathbb{R}^{m}$ is defined as:

$$d_{F}(\gamma_{1},\gamma_{2})=\inf_{\alpha,\beta}\max_{t\in[0,1]}\parallel\gamma_{1}(\alpha(t))-\gamma_{2}(\beta(t))\parallel$$

where $\alpha,\beta:[0,1]\to[0,1]$ range over all continuous non-decreasing reparametrizations. For $D_{\text{dev}}$, $\gamma_{1}=\gamma_{\text{elim}}$ is the post-peak elimination-phase segment of the delay-embedded trajectory, and $\gamma_{2}=L$ is the straight line connecting the start and end points of $\gamma_{\text{elim}}$.

Computation was performed using a discrete approximation with dynamic programming. Let $\gamma_{1}$ be discretized into $n_{1}$ points $\mathbf{p}_{1},\ldots,\mathbf{p}_{n_{1}}$ and $\gamma_{2}$ into $n_{2}$ points $\mathbf{q}_{1},\ldots,\mathbf{q}_{n_{2}}$. A dynamic programming table $D[i,j]$ is initialized with $D[1,1]=\parallel\mathbf{p}_{1}-\mathbf{q}_{1}\parallel$ and filled recursively for $i=2,\ldots,n_{1}$ and $j=2,\ldots,n_{2}$:

$$D[i,j]=\max\left( \parallel\mathbf{p}_{i}-\mathbf{q}_{j}\parallel,\min\left( D[i-1,j],D[i,j-1],D[i-1,j-1] \right) \right)$$

The discrete Fréchet distance is $D[n_{1},n_{2}]$, which converges to the continuous Fréchet distance as the discretization is refined.

**Data and Code Availability**

All simulation code used to generate the results in this paper, including the PTI vector extraction pipeline implemented in Python (version 3.9) using Ripser and GUDHI for persistent homology computation, is available at <https://github.com/yxgu2353/TopoPK>. The complete simulated datasets and PTI output tables are included in this Supplementary Information document. The pipeline generates all PTI parameters in a fully automated workflow requiring only the concentration–time curve and the embedding parameters ($m$, $\tau$) as inputs.
